# The Role of Temporal Factors in Processing Rapid Serial Visual Presentations

**DOI:** 10.64898/2025.12.15.694535

**Authors:** Cameron K. Phan, Imogen Breen, David Alais, Reuben Rideaux

**Affiliations:** School of Psychology, The University of Sydney, Sydney, NSW 2006, Australia; Cardiff University, Cardiff CF10 3AT, United Kingdom; Queensland Brain Institute, The University of Queensland, Brisbane, QLD 4067, Australia

**Keywords:** Acknowledgements: This work was supported by an Australian Research Council (ARC, Australia) Discovery Project Grant (DP250100118) and a National Health and Medical Research Council (NHMRC, Australia) Investigator Grant (2026318) awarded to RR

## Abstract

Sequentially presented visual stimuli are often employed to investigate the temporal limits of our visual perception and attention. Due to the finite nature of the brains’ processing capacity, experimental designs attempt to increase temporal separation within series to mitigate potential neural interference between stimuli. This is achieved through shortening of stimulus durations and lengthening of interstimulus intervals. Across two experiments, we parametrically varied these parameters for a serially presented checkerboard wedge stimulus and used inverted encoding modelling to assess the fidelity of the neural representation of spatial location. We found longer interstimulus intervals and shorter stimulus durations produced the most reliable decoding. Varying interstimulus interval altered the fidelity of neural representations via inter-item interference. By contrast, the influence of stimulus duration seemed largely analytic; in a fixed duration block, shorter stimulus duration enabled more presentations, which led to increased decoding accuracy. Distinct anticipatory and reactionary neural events were observed for stimuli presented in triplet sequences, which seemed consistent with forwards and backwards visual masking. The absence of an anticipatory event was observed when backward masking was stronger while a reactionary event was present for every stimulus subjected to forward masking.

## INTRODUCTION

Perception and cognition are frequently studied using sequentially presented visual stimuli (Keysers & Perrett, 2002; Mack et al., 2008; Maguire & Howe, 2016; Potter, 2012; Potter et al., 2014; Reeves, 1996; Robinson et al., 2019; Schiller & Smith, 1965). Research on the temporal limits of vision and attention has often employed the use of rapid presentation streams in what is known as Rapid Serial Visual Presentation (RSVP; Maguire & Howe, 2016; Potter, 2012; Potter et al., 2014). For each stimulus, a transient neural representation of its features is elicited. As stimuli are presented at higher frequencies, the brain is increasingly required to represent multiple stimuli simultaneously. There is a finite capacity for sensory processing in the brain, which places limits on this concurrent representation (Dux & Marois, 2009; Keysers & Perrett, 2002; Macknik & Martinez-Conde, 2008; Zivony & Lamy, 2022). While rapid serial presentation is a common approach in sensory experiments, the interaction between the temporal dynamics of stimulus presentation and neural limitations remains uncertain.

Behavioural work has shown that the accuracy of observers’ responses to visual stimuli is influenced by the temporal dynamics of presentation, such that more errors are made when stimuli are presented rapidly, which has been interpreted as approaching the temporal limits of visual perception (Maguire & Howe, 2016; Reeves, 1996). For example, when observers are exposed to two visual flashes in succession, they are more likely to report only perceiving a single flash when the non-presentation interval, the interstimulus interval (ISI), is shorter (Deodato & Melcher, 2024; Pearson & Tong, 1968; Reeves, 1996; Samaha & Postle, 2015). This fusion effect is typically found for ISIs under 50 ms and has been used to assign a limit on the visual system’s capacity to sample new information (Deodato & Melcher, 2024; Samaha & Postle, 2015). The minimum ISI required for an observer to report separate flashes appears to correlate with observers’ closed-eye alpha frequencies, a frequency range typically associated with inhibition and sampling, but not with beta or theta frequencies (Deodato & Melcher, 2024).

Another set of behavioural phenomena associated with serial visual presentations are masking effects (Macknik & Martinez-Conde, 2008; Schiller & Smith, 1965). Masking occurs when the visibility of a stimulus is reduced because another stimulus, presented shortly before (forward masking) or after (backward masking), raises the threshold required to perceive it. Both forward and backward masking strength decreases (i.e., result in lower perceptual thresholds of the target stimulus) as the stimulus onset asynchrony (SOA) between mask and target stimuli increases, reaching a minimum at ∼150 ms (Macknik & Martinez-Conde, 2008; Schiller & Smith, 1965). As opposed to the resolution limit of processing proposed for the fusion effect, interference between the overlapping processing of sequential stimuli is thought to reduce the available information to the brain, effectively reducing the signal-to-noise ratio (Eriksen & Schultz, 1978; Keysers & Perrett, 2002; Macknik & Martinez-Conde, 2008). Supporting evidence from neurophysiological work shows decreased amplitude in the average peak activity and decreased neuronal spiking rates (Noguchi & Kakigi, 2005; Rolls et al., 1999). Both fusion and visual masking effects are typically considered products of low-level perceptual processing limits that operate in the primary visual cortex (V1) at short timescales (Deodato & Melcher, 2024; Green et al., 2005; Rieger et al., 2005; Samaha & Postle, 2015). Unlike the long sequences associated with RSVP, these effects are often studied by presenting pairs of stimuli (Deodato & Melcher, 2024; Macknik & Martinez-Conde, 2008; Pearson & Tong, 1968; Samaha & Postle, 2015; Schiller & Smith, 1965).

The attentional blink is a phenomenon widely studied with RSVP (Dux & Marois, 2009; Shapiro et al., 1997; Zivony & Lamy, 2022), where observers tasked to detect two targets fail to do so for the second target when it is presented within 500 ms after the first. Like forward masking, the cause of the attentional blink was initially considered to be a result of interference in forming the perceptual representation of the second target due to the processing of the first (Dux & Marois, 2009). However, this explanation is challenged by the finding that the attentional blink occurs for targets that are visually distinct (thus, unlikely to produce interference) and is dependent on task demands. The interference explanation is further challenged by a lack of correlation between the attentional blink effect and the strength of P1 evoked response potentials (ERP) in electroencephalography (EEG) recordings, an ERP associated with the onset of visual processing (Eimer, 1996, 2014; Polich, 2007; Zivony & Lamy, 2022). Rather, the amplitude of later ERP components associated with attentional engagement and working memory were found to correlate with attentional blink strength. Therefore, instead of the low-level perceptual limits that drive fusion and visual masking, the attentional blink appears to be a product of later attentional or memory capacity limits.

In the past decade, multivariate decoding approaches have increasingly been used to characterise neural responses elicited by rapidly presented stimuli, recorded using EEG (Grootswagers et al., 2019; Robinson et al., 2019). The reliability of decoding reflects the distinctiveness of neural patterns, and differences in decoding accuracy produced by experimental manipulations can provide insight into the neural mechanisms that both serve and limit rapid sensory processing. For instance, the interference associated with fusion, masking, and the attentional blink, would be expected to manifest as reduced decoding accuracy, the timing of which can provide evidence for the neural mechanisms that are impacted.

Traditionally, experiments employing the RSVP paradigm have sought to balance minimising the potential interference between processing of stimuli and maximising the number of presentations. The implication is that as temporal separation of stimuli increases, interference is less likely to occur. Consequently, within a given experimental session, the number of stimulus presentations is inversely proportional to the SOA of the stimuli. Electrophysiological work employing multivariate decoding of EEG recordings has shown supporting evidence of this interference in the neural representations (Grootswagers et al., 2019; Robinson et al., 2019). In particular, longer SOAs are associated with higher decoding accuracy. Given that SOAs are comprised of two components, the stimulus duration and the ISI, either one or both could have contributed to these findings. Longer pulse durations produce lower flash fusion thresholds, attributed to the decreased persistence of the first flash and its interference with the second (Purcell & Stewart, 1971). Similarly, detection of targets presented in RSVP is modestly improved when stimuli are presented for longer (Maguire & Howe, 2016; Potter et al., 2014). However, this benefit does not transfer to the attentional blink, i.e., stimulus duration does not influence the magnitude of the effect (Dux & Marois, 2009), nor does duration seem to affect the decoding accuracy of stimuli presented in RSVP (Robinson et al., 2019).

In contrast to stimulus duration, longer ISIs typically increase both behavioural performance and EEG decoding accuracy. In RSVP studies, observers’ target detection performance improves when ISIs are introduced or increased (Intraub, 1980; Nieuwenstein et al., 2009; Potter et al., 2004). Similarly, the attentional blink is reduced when a blank period is presented between target stimuli, such that the likelihood of detecting of the second target is increased (Nieuwenstein et al., 2009). This is also seen for visual masking, where extending the ISI increases detection performance of stimuli presented either after or before the mask (Maguire & Howe, 2016; Potter et al., 2014; Schiller & Smith, 1965). These behavioural effects are mirrored by those observed in neuroimaging work; introduction of an ISI between stimuli increases the decoding accuracy of stimulus features (Robinson et al., 2019). Whether the effect of ISI on decoding accuracy is graded, like behavioural performance effects, remains unknown.

To investigate the influence of temporal dynamics on the processing of serially presented visual stimuli, here we used multivariate analysis to decode EEG recordings of responses to wedge stimuli presented at different polar angles. Presenting the stimuli at different spatial location minimises the potential of spatial masking that may occur for stimuli presented in the same or overlapping spatial locations (Enns, 2004; Herzog, 2008). We separately manipulated both stimulus duration and ISI to test their effects on serial processing. In Experiment 1, we determined how these factors influence the decoding accuracy of spatial location. We found that when controlling for the number of presentations, ISI, but not stimulus direction, influenced overall decoding accuracy. In Experiment 2, we tested whether the effect of ISI on decoding accuracy *i*) reflected perceptual performance, and *ii*) was due to forwards or backwards masking/interference. The results showed the difference in decoding accuracy was mirrored by perceptual recall of stimuli and suggested that both forwards and backwards masking influenced the stimulus representation. Additionally, we found evidence of anticipatory and reactive neural re-activation of the stimulus representation, in response to the presentation of subsequent stimuli. These findings provide a roadmap for optimizing the temporal dynamics of RSVP experiments. Further, the findings provide cross-validation for neural decoding accuracy as an index of perceptual fidelity, highlight the presence of extra-spatial masking (forwards and backwards), and reveal neural mechanisms that may serve to preserve the identity of previous stimuli through an anticipatory re-activation prior to predict incoming stimulation.

## METHODS

### Participants

There were 21 (mean ±standard deviation age, 25.5 ±6.3, 14 female) and 12 (mean ±standard deviation age, 26.0 ±4.2, 9 female) healthy human participants for Experiments 1 and 2, respectively. Both samples were recruited from The University of Sydney. Neither experiment had any exclusions. Each participant provided informed consent before commencing and had normal or corrected-to-normal vision (assessed using a standard Snellen eye chart). The study protocol was approved by The University of Sydney Human Research Ethics Committee (2023/HE000072). Participants received either payment or course credit in return for their time.

### Apparatus and Stimuli

Both experiments were conducted in a dark, acoustically and electromagnetically shielded room. The stimuli were presented on a 24-inch ASUS VG248QE Gaming Monitor (ASUS, Taipei, Taiwan) with 1920 x 1080 resolution and a refresh rate of 144 Hz. The monitor was calibrated using a SpyderX Pro (Datacolor, Lawrenceville, NJ). Viewing distance was maintained at 70 cm using a forehead and chinrest, meaning the screen subtended 41.47° x 23.33° (each pixel 1.3’ x 1.3’). Stimuli were generated in MATLAB v2020a (The MathWorks, Inc., Matick, MA) using Psychophysics Toolbox (Brainard, 1997; Pelli, 1997) v3.0.18.13 (see http://psychtoolbox.org/). The visual stimuli consisted of black and white checkered wedges subtending 22.5° in width and extended 6° of visual angle from a central fixation dot (0.2° visual angle radius). The wedge stimuli were presented at polar angles that were randomly sampled between 0° and 360°. Continuous EEG was recorded on a 64-channel BrainVision ActiCap system (Brain Products GmbH).

### Procedure

In Experiment 1, participants were instructed to maintain fixation on the centrally presented dot. Participants completed two 25-min blocks with a break between blocks that was limited to a maximum of five min. The blocks were comprised of one-min sequences for each of the 25 stimulus conditions (**Fig. 1a**), totalling 25 min. The 25 conditions were the product of a full factorial design of five stimulus durations and five ISIs, each factor ranged from 50 to 850 ms (in 200 ms steps). As such, the stimulus presentation frequency ranged from 0.59 to 10 Hz and the stimulus presentations per condition ranged from 70 to 1200 in total. The order of the 25 different condition sequences was randomly interleaved within each block. After each 1-min sequence, the task for half the participants was to indicate whether there were more wedges presented on the left or right, while the other half indicated whether there were more above or below fixation. Feedback was provided.

**Figure 1.**
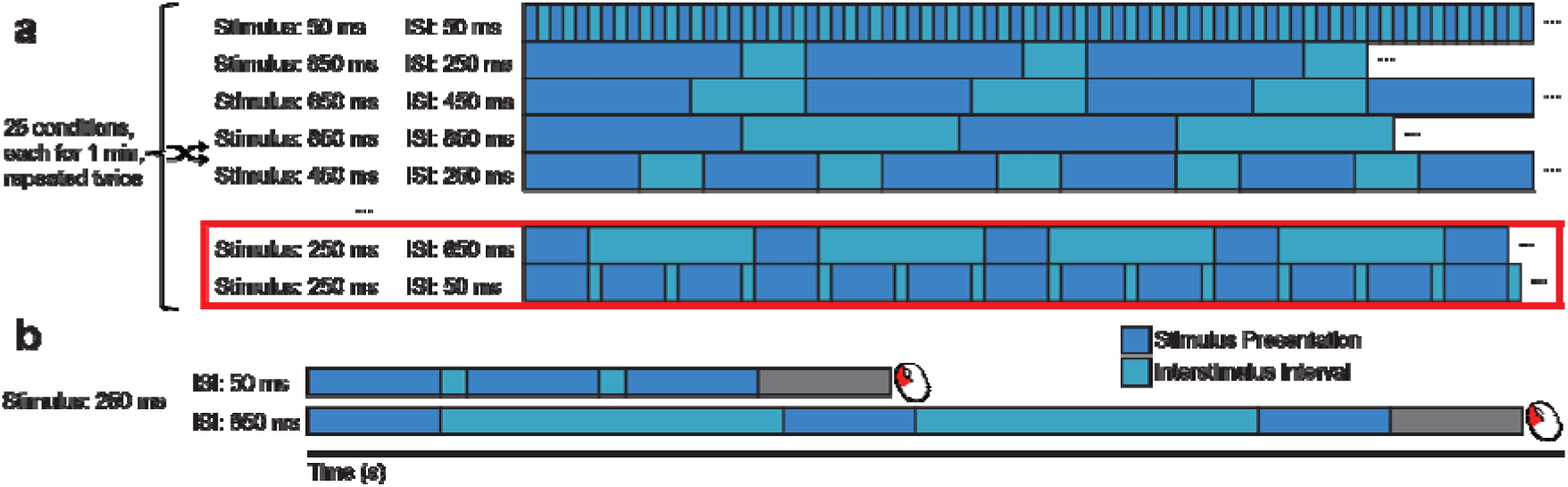
Schematic of temporal configurations of presentation sequences. **a**) Display of the 25 stimulus conditions resulting from the 5 (stimulus duration) × 5 (interstimulus interval) design employed in Experiment 1. The conditions outlined in red were selected for Experiment 2, the sequence structure of which is shown in (**b**).

In Experiment 2, participants were instructed to maintain fixation on the centrally presented dot. Each participant completed three runs. Each run was comprised of 10 blocks, each of which contained 20 sequences. Sequences consisted of three stimulus presentations. The stimuli were presented for 250 ms followed by the ISI of the current block. This resulted in 900-ms sequences for the 50 ms ISI condition and 2700-ms sequences for the 650 ms ISI condition (**Fig. 1b**). After each sequence, the participant’s task was to reproduce the position of the second wedge stimulus by using the mouse to rotate a checkered wedge to the target location.

### EEG

EEG recordings were digitized at a 1024 Hz sampling rate using a 24-bit A/D conversion. The 64 Ag/AgCl electrodes were arranged on the scalp aligning with the 10–20 international standard (Oostenveld & Praamstra, 2001), via a nylon cap. All electrodes were referenced to FCz. Offline EEG pre-processing was performed using EEGLAB v2025.0.0 (Delorme & Makeig, 2004). The data were initially down-sampled to 256 Hz and subjected to a 0.1 Hz high-pass filter to remove slow baseline drifts and a 45.0 Hz low-pass filter to remove high-frequency noise/artifacts. Data were then re-referenced to the common average before being epoched into segments around each stimulus. The epochs extended from -0.2 s to 2 s from the onset of stimulus presentation for Experiment 1 and extended from -0.2 s to 3.3 s from the onset of stimulus presentation for Experiment 2. As such, the epochs for certain conditions contained ERPs produced from multiple stimuli. However, stimuli were deliberately randomly sampled, such these additional signals could not provide any information that could be used to decode the current stimulus. For each epoch, the mean EEG sensor activity from the pre-stimulus period (i.e., -0.2 s to 0 s from onset of stimulus presentation) was subtracted from the remainder of the epoch data as a means of baselining.

### Neural Decoding

An inverted modelling approach was applied to the pre-processed EEG data to reconstruct the polar angle location of the checkered wedge stimuli (Brouwer & Heeger, 2011). A forward model was nominated that described the measured activity in the EEG sensors given the polar angle of the presented wedge stimulus. The forward model was then used to obtain the inverse model that described the transformation from EEG sensor activity to the polar angle of the presented wedge stimulus. For the main decoding analyses, the forward and inverse models were obtained using a ten-fold cross-validation approach in which 90% of the data were used to obtain the inverse model on which the remaining 10% were decoded.

The forward model consists of five hypothetical channels (Brouwer & Heeger, 2009) evenly distributed between 0° and 360° (i.e., the first channel was centred at 36° with a width of 72°). Each channel consisted of a half-cycle sinusoid raised to the fourth power to effectively emulate a half-wave rectified sinusoid, albeit with a smooth transition to 0 values. The tuning curve of any polar angle could then be expressed as a weighted sum of the five channels. This can be expressed as the following linear equation:

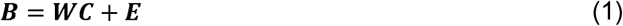

where ***B*** (*m* sensors × *n* presentations) is the pre-processed EEG electrode data, ***W*** (*m* sensors × 5 channels) is the weight matrix used to transform the EEG data to wedge stimulus polar angle, ***C*** is the hypothesised channel activities corresponding to the polar angle of the presented wedge stimulus, and ***E*** indicates the residual errors.

For the inverse model, we fitted for weights that when applied to the pre-processed EEG electrode data reconstructed the underlying channel activities with the least residual error. To optimise the inverse model for the EEG data, we took the noise covariance into account as a means of combating the high correlations between neighbouring sensor activity (Kok et al., 2017; Mostert et al., 2015; Rideaux, 2024). ***B*** and ***C*** were demeaned such that their average over presentations equalled zero for each sensor and channel, respectively. The inverse model was then estimated using either a subset selected through cross-fold validation or all the data in one condition. The hypothetical responses of each of the five channels were calculated from the training data, resulting in the response row vector ***c**_train,i_* of length *n_train_* presentations for each channel *i*. The weights on the sensors ***w**_i_* were then obtained through least squares estimation for each channel:

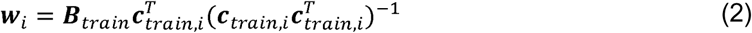

where ***B**_train_* is the (*m* sensors × *n_train_* presentations) training portion of the pre-processed EEG electrode data. The optimal spatial filter vi to recover the activity of the ith channel was obtained as follows (Mostert et al., 2015):

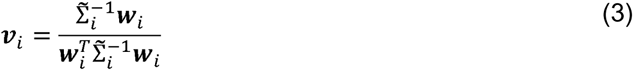

where Σ̃_*i*_ is the regularised covariance matrix for channel *i*. Incorporating the noise covariance in the filter estimation leads to the suppression of noise that arises from correlations between sensors. The noise covariance was estimated as follows:

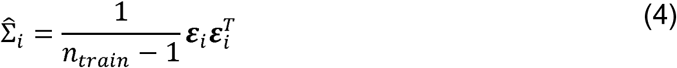

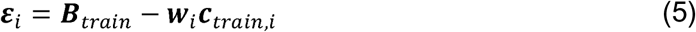

where *n_train_* is the number of training presentations. For optimal noise suppression, we improved this estimation by means of regularisation by shrinkage using the analytically determined optimal shrinkage parameter (Mostert et al., 2015), yielding the regularised covariance matrix Σ̃_*i*_.

For each presentation, we decoded polar angle location by converting the channel responses to polar angles:

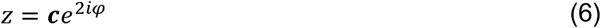

and calculating the estimated angle:

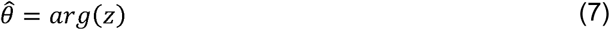

where is a vector of channel responses and φ is the vector of angles at which the channels are centred. From this estimate we calculated accuracy. Accuracy represented the similarity of the decoded polar angle to the presented polar angle (Kok et al., 2017), and was expressed by the following equation:

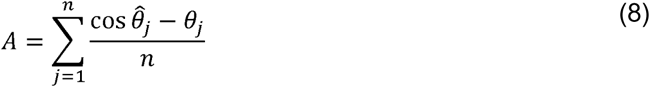

where accuracy *A* is the mean of the cosine of the difference between the decoded polar angle *θ̂*_*j*_ and the corresponding presented polar angle *θ*_*j*_ across n presentations, providing a performance measure on a normalised -1 to 1 scale, where positive differences indicated a decoded angle counterclockwise to the presented angle.

### Statistical Analysis

Statistical analyses were performed in MATLAB v2025a, CircStat Toolbox v2012a (Berens, 2009), and using Mixture Modelling from the Analogue Report Toolbox (Bays et al., 2009; Schneegans & Bays, 2016). For each condition and time point, the *t*-statistic was calculated as a proxy for effect size of decoding accuracy (**Fig. S1 & S2**). Contiguous time points with significant values were demarcated into clusters. To simulate a null distribution of the maximum summed cluster values, we permuted (*n* = 5000) the sign labels of the decoding accuracy data, for single, respectively. A 95% percentile threshold value was derived from this null distribution. Clusters with summed absolute *t*-statistics smaller than this threshold were considered not significant.

For the determination of the significance of the main effects, a similar cluster correction was applied to repeated-measures ANOVA *F*-statistics. The condition labels of each main effect were permuted (*n =* 5000) for each participant to produce null datasets, which had repeated-measures ANOVAs performed and thereby produced a null distribution of *F*-statistics. Clusters of significant *F*-statistics were summed in the data and compared to the 95% percentile of the null distribution. Clusters with summed *F*-statistics less than this threshold were deemed not significant.

As a consequence of matching the duration of blocks across conditions in Experiment 1, each condition had an unequal number of stimuli presentations. The conditions in which there were more presentations may have benefitted from better decoding accuracy due to the increased number of presentations available to establish a reliable model of the neural representation. To assess the influence of unequal presentations, we repeated the neural decoding and analyses after matching the number of presentations across conditions to the condition with the fewest presentations. To match the number of presentations, we randomly selected without replacement the stimuli to analyse within each condition for each participant.

For the behavioural responses in Experiment 2, mixture modelling (Bays et al., 2009; Schneegans & Bays, 2016) was used to extract the probability of participants’ responses being derived from recalling either of the nontarget stimuli (i.e., first or third in sequence) and their probability of producing random guess response from the overall rate of errors they made. Additionally, the kappa parameter of the von Mises distributions of the participants’ responses was calculated as a means of determining the precision of their responses. Shapiro-Wilk tests for normality were conducted on each of these behavioural performance measures for each interstimulus interval condition. Where normality was violated, nonparametric Wilcoxon signed-rank tests were conducted between the interstimulus intervals, otherwise, one-way repeated-measures ANOVAs were employed instead.

## RESULTS

Across all conditions, the decoding accuracy of polar angle increased sharply ∼60 ms post stimulus onset (**Fig. 2**). Reliable decoding accuracy persisted until ∼400 ms following stimulus onset, despite new stimuli continuing to arrive within this 400-ms period in most of the conditions.

**Figure 2.**
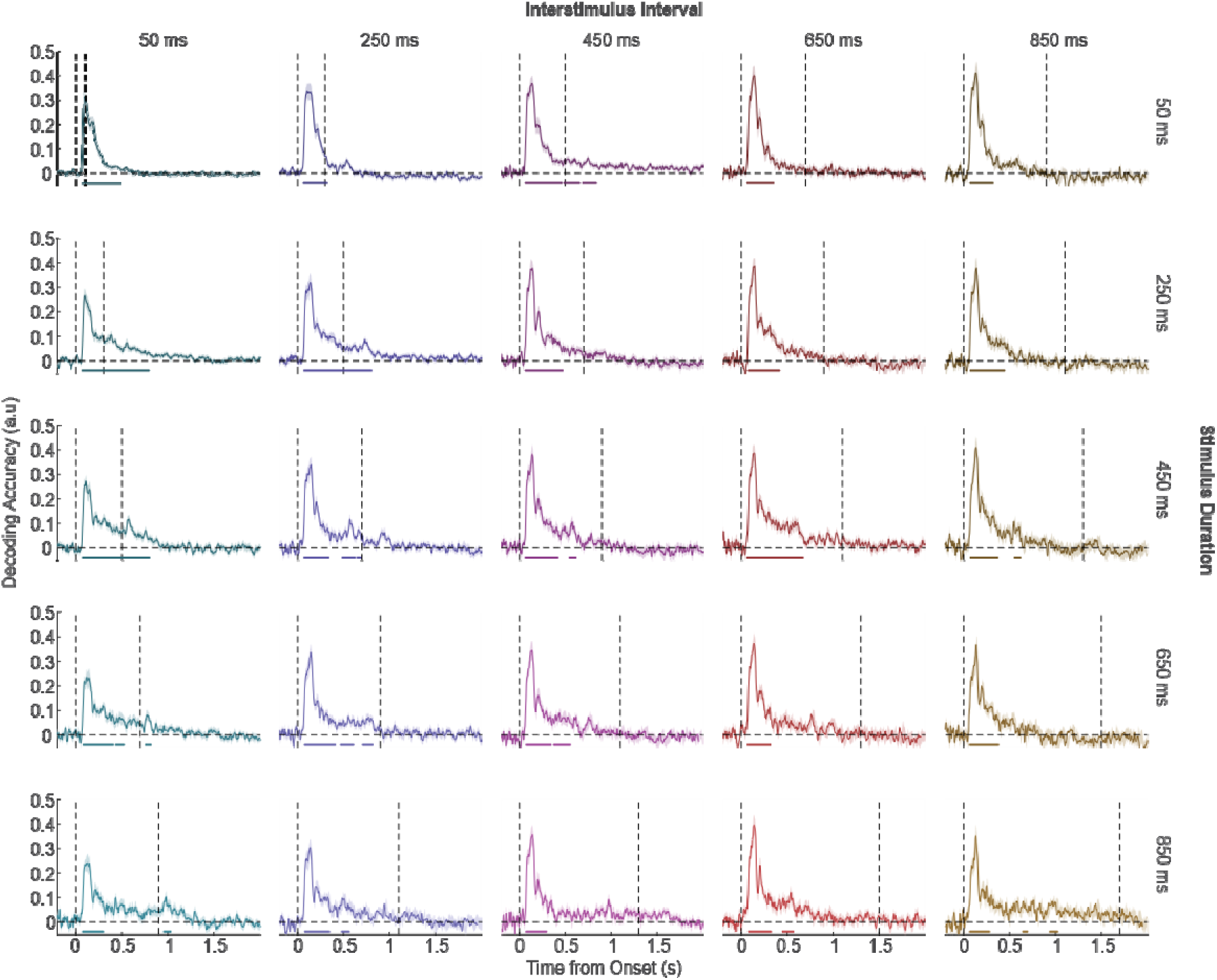
Decoding accuracy for all stimulus conditions. Average decoding accuracy for all 25 conditions of Experiment 1 from -0.2 s to 2 s from stimulus onset. Condition labels are indicated by the column and row headings (right-hand side). Vertical dashed lines indicate the onset of the target (epoched) stimulus and the subsequent stimulus. Shaded regions indicate ±1 SEM and coloured horizontal bars show cluster-corrected periods of significance.

To assess the influence of stimulus duration on the neural representation of stimulus features, we averaged decoding accuracy across ISI conditions and compared performance between duration conditions (**Fig. 3a**). We found multiple periods in which decoding accuracy was significantly different between stimulus duration conditions. During early periods (<200 ms), there was a clear ordering of conditions indicating better decoding accuracy for shorter durations. However, the order was less clear for the later periods. We performed the same analysis for ISI and found two (early) periods in which decoding accuracy was significantly different between conditions (**Fig. 3b**). In contrast to stimulus duration, the ordering of conditions for ISI during these periods showed a benefit for longer ISIs. Indeed, these early influences of stimulus duration and ISI can be observed when decoding accuracy is temporally averaged over the initial stages of stimulus processing (0.05 to 0.3 s; **Fig. 3c**).

**Figure 3.**
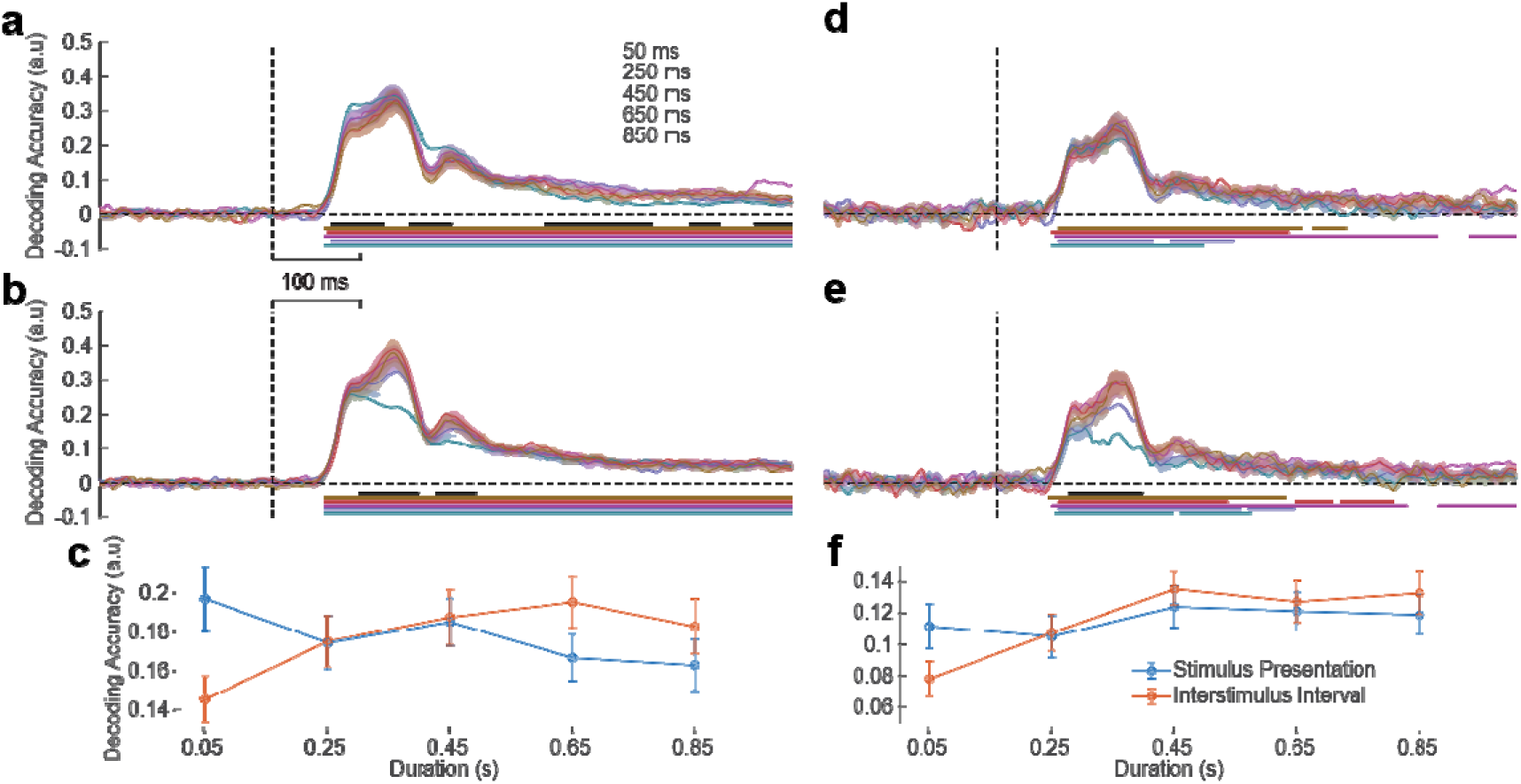
The influence of stimulus duration and ISI on neural representations. **a**) Decoding accuracy for the five stimulus durations, averaged over interstimulus interval. **b**) Same as (**a**), but for the five interstimulus intervals, averaged over stimulus duration. **c**) The results from (**a** & **b**) temporally averaged from 0.05 s to 0.3 s from stimulus onset. The number of stimulus presentations within each condition were matched for the one-min duration of the blocks, resulting in a range from 70 to 1200 stimulus presentations. (**d**-**f**) Same as (**a**-**c**), but matching the number of presentations (70) across conditions. Horizontal bars indicate cluster-corrected periods of significance, where black bars show differences between conditions and coloured bars show differences between specific conditions and chance-level decoding. The shaded regions in (**a**) and (**b**) and the error bars in (**c**) represent ±1 SEM.

When the number of presentations were matched, we no longer found any significant differences between stimulus duration conditions, averaged across ISI (**Fig. 3d**). Averaged over stimulus duration, the first of the two periods of significant differences between ISI conditions persisted and demonstrated the same benefit of longer ISIs for decoding accuracy as when block duration was matched (**Fig. 3e**). These changes in the early influence of stimulus duration and ISI can be seen when temporally averaged over the initial stages of stimulus processing (**Fig. 3f**).

The results of Experiment 1 indicate that for sequential presentation of visual stimuli, the ISI influences the decoding accuracy of stimulus features. Higher accuracy, indicative of more distinguishable and reliable patterns of neural activity, was found for stimuli presented with longer ISIs, even when the number of stimulus presentations used to train the decoder is matched. This result is in line with the findings from Robinson et al. (2019), where non-zero ISIs resulted in greater decoding accuracy of RSVP stimuli, and consistent with the improved memory of stimuli presented with longer ISIs found in behavioural research (Intraub, 1980; Nieuwenstein et al., 2009; Potter et al., 2004). While our decoding results are consistent with previous behavioural work, it remains uncertain whether these neural and behavioural phenomena are associated, that is, whether the differences in decoding accuracy reflect differences in perceptual fidelity. For instance, reduced decoding accuracy may reflect genuine neural interference between stimuli, reducing perceptual fidelity. However, given the spatially coarse sampling of EEG recordings, it may simply reflect interference between neural responses at the sensor level. Further, while previous work has suggested that masking likely explains the influence of ISI on decoding accuracy, our stimuli were largely presented in spatially non-overlapping locations, which challenges the standard explanation of spatial masking. Adding further uncertainty to the masking explanation is that in previous studies and here, both backwards and forwards masking occur for all stimuli, except the first and last in a sequence. Thus, it remains unclear whether one or both types of masking are responsible for influence of ISI.

To address these questions, we ran a second experiment that was similar to the first, but with shorter stimulus presentation sequences of three stimuli. Within the three stimuli sequences presented, the first and third were subjected only to backwards and forwards masking, respectively, whilst the second is subject to both. Thus, differences in decoding accuracy between the first and third stimuli should provide an indication of the relative strength of the directional masking effects. Previous behavioural work has shown that forward masking is stronger than backward masking, producing greater interference with the processing of the stimulus following the masking stimulus (Schiller & Smith, 1965). Overall, masking should decrease with ISI as the interference between stimulus processing increases with temporal proximity. In the interest of maximising potential masking effects, we selected the temporal configurations that had the greatest pairwise difference in decoding accuracy in Experiment 1, when the number of stimulus presentation was matched. Specifically, we used 250 ms stimulus duration for both with 50 and 650 ms ISI, respectively. Based on the results of Experiment 1, we expected better decoding accuracy for stimuli presented with a longer ISI, reflecting diminished masking interference.

Additionally, to test whether the reduced decoding accuracy associated with shorter ISIs reflects diminished perceptual fidelity, participants in Experiment 2 were tasked with reproducing of the location of the second stimulus after each sequence. If the differences in decoding accuracy were primarily driven by sensor-level interference, the behavioural responses should not show any significant improvement for target stimuli presented with the longer (650 ms) ISIs, as would be expected with reduced interference between neural representations.

Participants’ behavioural responses in Experiment 2 could have consisted of correct replications of the second stimulus, incorrect replications of either non-target (first or third) stimuli, or guess responses. As such, we applied mixture modelling to participants’ responses to estimate the precision of their target responses, the probability of their responses being made to non-target stimuli (swap errors), and the probability that their responses were guesses. Shapiro-Wilk tests revealed that normality was violated for reproduction precision (50 ms ISI: *W* = .752, *p* = .004; 650 ms ISI: *W* = .763, *p* = .005) and guess rates (50 ms ISI: *W* = .468, *p* < .001; 650 ms ISI: *W* = .850, *p* = .036). In both these cases, the non-parametric Wilcoxon signed-rank test was used to determine the significance of differences between ISIs. We found that reproduction precision was significantly better for stimuli presented in sequences with 650 ms ISIs than those with 50 ms intervals (*W* = 0, *p* < .001 ; **Fig. 4a**). By contrast, we found no difference in swap rate between conditions (*F*_1, 11_ = 3.15, *p* = .104, *BF_10_* = 1.307; **Fig. 4b**). For guess rate, we found a higher rate when using a 650 ms ISI than one of 50 ms (*W* = 1, *p* = .017; **Fig. 4c**).

**Figure 4.**
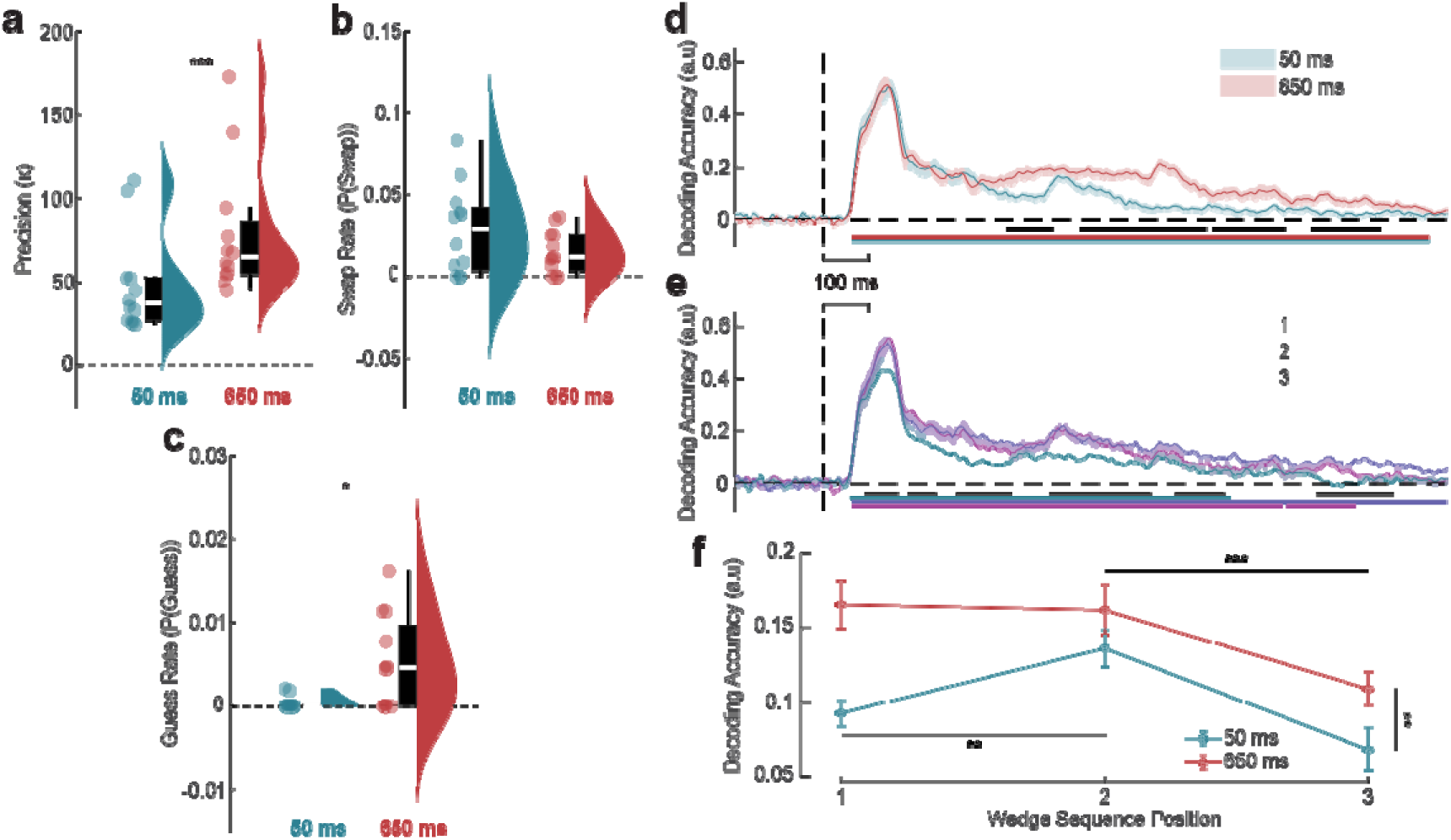
Influence of ISI and order on stimulus reproductions and neural representations. **a-c)** Participants’ (**a**) reproduction precision, (**b**) swap rate, and (**c**) guess rate in response to sequences of stimuli with ISIs of 50 and 650 ms. **d**) Decoding accuracy for the two ISIs, averaged across sequence position. **e**) Same as (**d**), but for sequence position, averaged over ISI. **f**) The results of (**d** & **e**) temporally averaged from 0.05 s to 1.4 s from stimulus onset. Horizontal bars indicate cluster-corrected periods of significance, where black bars show differences between conditions and coloured bars show differences between specific conditions and chance-level decoding. The shaded regions in (**d**) and (**e**) and the error bars in (**f**) represent ±1 SEM. \**p* < .05, \*\**p* < .01, \*\*\**p* < .001.

Decoding accuracy of stimulus location across all stimulus conditions increased sharply ∼60 ms post stimulus onset and persisted until ∼1200 ms (**Fig. 4d, e**). We first averaged decoding accuracy across sequence position and found better performance for 650 than 50 ms ISIs (**Fig. 4d**). In contrast with Experiment 1, the periods of significant difference occurred later (>350 ms). We averaged across ISI to test the effect of sequence position and found periods of significant difference in decoding accuracy spread across the epoch (**Fig. 4e**). For the first four periods (<700 ms), the difference was driven by significantly lower decoding accuracy for the third stimulus. For the later periods, the difference was primarily due to the second stimulus having higher accuracy. For the second of these periods (∼1100 ms), an ordered difference manifested in the first stimulus having significantly higher decoding accuracy than the third.

Averaged across all time points post stimulus onset (**Fig. 4f**), there was a significant interaction between ISI and sequence position on decoding accuracy (*F*_2, 11_ = 3.696, *p* = .041, *BF_10_* = 1.821). The second stimulus was decoded with significantly greater accuracy than the first when the ISI was 50 ms (*t*_11_ = 3.614, *p* = .004), but not 650 ms (*t*_11_ = 0.197, *p* = .848). For both ISI conditions, the decoding accuracy of the second stimulus was significantly greater than the third (50 ms ISI: *t*_11_ = 3.354, *p* = .006; 650 ms ISI: *t*_11_ = 3.130, *p* = .010; *t*_11_ = 4.924, *p* < .001). This difference between the second and third stimulus was not significantly different between ISIs (*t*_11_ = 0.528, *p* = .608).

To further explore the influence of ISI on the neural representation of the stimulus, we calculated decoding accuracy separately for each ISI and sequence position (**Fig. 5**). We found the most differences between ISIs for the first stimulus. These emerged ∼100 ms after the onset of the second stimulus in 50 ms ISI sequences (∼400 ms). During this period, decoding accuracy decreased for the stimulus presented with 50 ms ISI while it remained stable in the 650 ms ISI condition. We found similar, yet less pronounced, pattern of differences for the second and third stimuli in the sequence.

**Figure 5.**
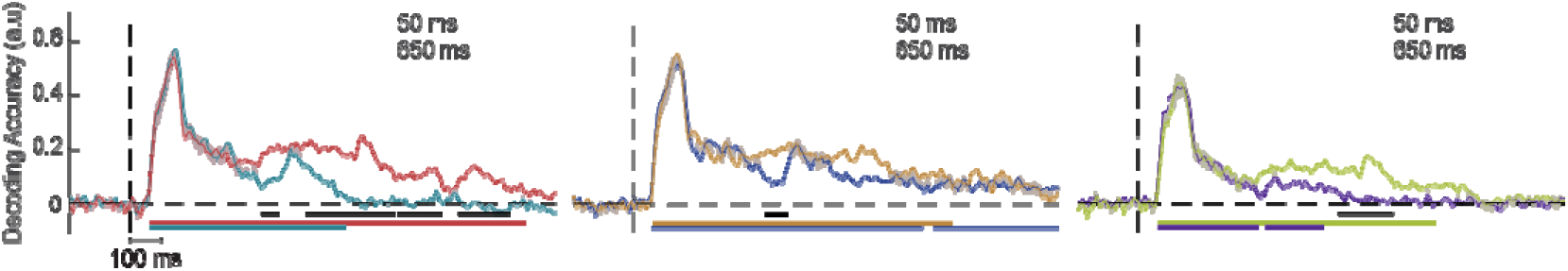
Influence of ISI on neural representations for each sequence position. Decoding accuracy for the interstimulus intervals for each of the three sequential positions (from left to right: first, second, and third stimulus). Horizontal bars indicate cluster-corrected periods of significance, where black bars show differences between interstimulus intervals and coloured bars show the difference between each interval condition and chance-level decoding. The shaded regions represent ±1 SEM.

To examine how subsequent stimuli influence decoding accuracy, we extended the epoch window around each stimulus to include the presentations of all three stimuli (−0.25 to 3.5 s around onset of the first stimulus; **Fig. 6**). From this analysis we observed three distinct phenomena. First, we found that for stimulus one and two in the sequence, there was a rise in decoding accuracy following the initial peak in the decoding accuracy of the second and third stimuli, respectively; as if the representation of the previous stimulus was re-activated by exposure to a new stimulus. Consistent with this explanation, no rise in decoding accuracy was observed at this temporal offset for the third stimulus in either ISI condition (**Fig. 6a, b**, vertical black dashed line). Second, for the target stimulus (2) in both ISI conditions, decoding accuracy underwent an additional increase after ∼50 ms from the offset of the third stimulus, which likely reflects preparation for the reproduction response of the target stimulus. Third, specifically for the stimuli presented with 650 ms ISI (**Fig. 6b**), decoding accuracy appears to spike followed by a steep decline immediately before the onset of subsequent stimuli. In this case, this pattern in decoding accuracy is found for the third stimuli at the time immediately preceding a hypothetical fourth stimulus (i.e., 900 ms after onset of the third stimulus or 2700 ms after the onset of the first stimulus; **Fig. 6b**, vertical black dashed line). Whereas the first phenomenon may reflect a reactive mechanism, this pattern of results likely reflects an anticipatory mechanism.

**Figure 6.**
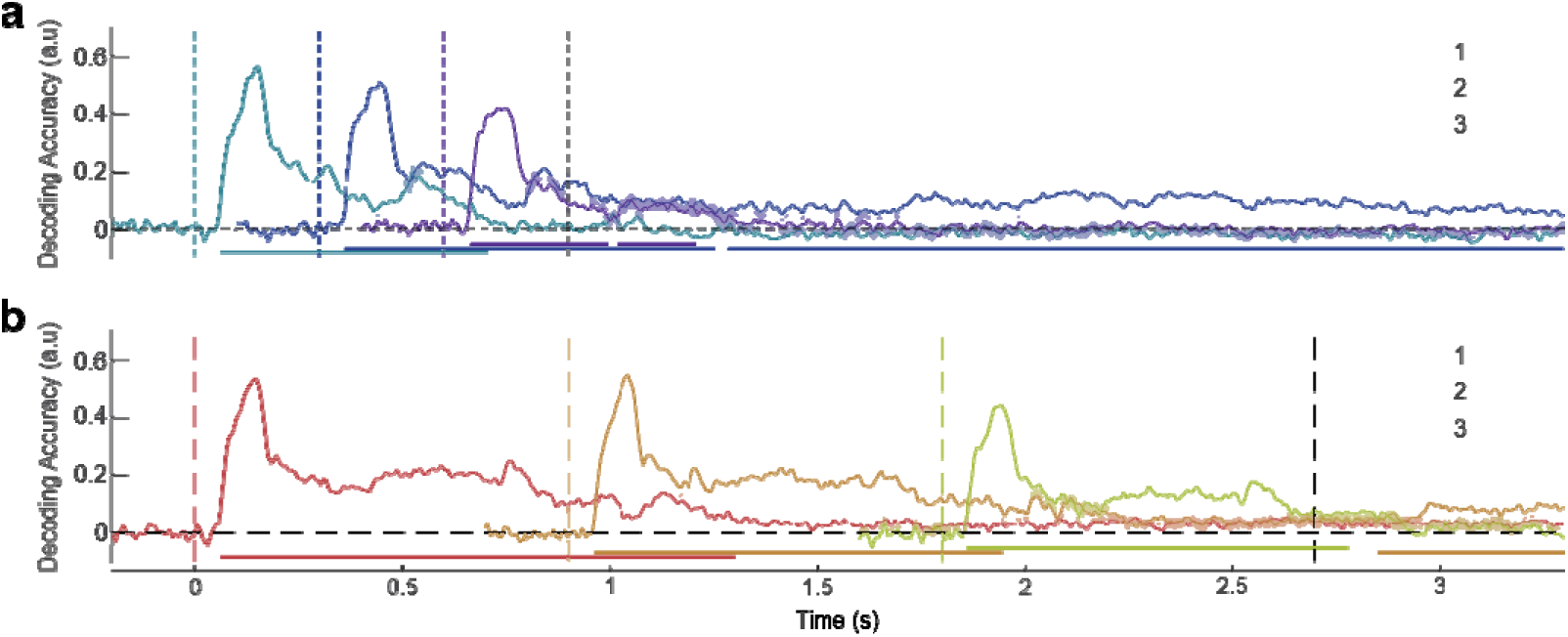
Time course of neural representations within sequences. Decoding accuracy of stimuli in each sequence position for (**a**) 50 ms and (**b**) 650 ms ISI. Vertical dashed lines indicate the onset of stimuli, where black is the onset of a hypothetical fourth stimulus and coloured lines correspond to the three presented stimuli. Coloured horizontal bars are for the difference between each condition and chance-level decoding. The shaded regions represent ±1 SEM.

## DISCUSSION

The current study sought to investigate the influence of stimulus duration and ISI on the neural representation of sequentially presented visual stimuli. In Experiment 1, we found that both stimulus duration and ISI influence how accurately stimuli can be decoded from EEG responses. In particular, shorter stimulus duration and longer ISI led to better decoding accuracy. However, the influence of stimulus duration seemed to be an outcome of the additional presentations associated with shorter duration when total presentation time was matched. Matching the number of presentations seemed to eliminate the influence of stimulus duration. This suggests that using brief stimulus presentations (e.g., 50 ms) provides the best outcome for decoding accuracy. However, it is possible that a longer stimulus duration would be optimal for decoding more complex features, such as object identity (Grootswagers et al., 2019; Retter et al., 2018; Robinson et al., 2019), as these take longer to process.

In contrast to the influence of stimulus duration, the benefit of longer ISIs persisted after controlling for stimulus presentations. Qualitative inspection of the data suggested that increasing ISIs improved decoding accuracy up until 650 ms, after which accuracy plateaued. This trend is somewhat similar to the detriment seen for secondary targets produced by attentional blink (Dux & Marois, 2009; Shapiro et al., 1997; Zivony & Lamy, 2022). However, previous work has shown that there is a short window (∼50 ms) after the presentation of the first target in which presentations of the second target will not induce an attentional blink effect (Dux & Marois, 2009; Hommel & Akyürek, 2005); whereas we found the lowest decoding accuracy for 50 ms ISI. Similarly, as the decoding accuracy of 50 ms ISI stimuli remained reliable, it is unlikely that the stimuli were perceptually fused as that would not have produced distinct neural representations (Deodato & Melcher, 2024; Samaha & Postle, 2015). Therefore, the most likely mechanism for the influence of ISI on decoding accuracy is masking. This interpretation is consistent with previous multivariate decoding studies (Robinson et al., 2019).

Experiment 2 sought to further investigate the influence of ISI on the neural representation of sequentially presented stimuli. Namely, whether changes in decoding accuracy reflected corresponding changes to perceptual fidelity, and the extent to which these effects were caused by forwards or backwards masking. Regarding the former, the behavioural results validated decoding accuracy as an index of perceptual fidelity; consistent with decoding performance, observers recalled items in the 50 ms ISI condition less precisely than those in the 650 ms condition.

Regarding masking, the results of Experiment 2 were less clear. If forwards and/or backwards masking determined the differences in decoding accuracy between the three stimuli presented in each sequence, there would be three possible patterns of results that could be expected. If only forwards or backwards masking occurred, we would expect the best decoding accuracy for the first or the third stimulus, respectively. If both occurred, we would expect the worst accuracy for the second stimulus, as it would be subject to both forwards and backwards masking. In contrast to these predictions, decoding accuracy of the second stimulus was either similar (650 ms ISI) or better (50 ms ISI) than that of the first and third. This is likely due to task demands; the second stimulus was task relevant, which may have engaged top-down attention mechanisms that mitigated the effect of masking. Considering this added influence of task demands, the results seem to indicate that forwards masking produced more interference than backwards masking, as decoding accuracy was lowest for the third stimulus in both ISI conditions. However, given the influence of task demands on decoding performance of the second stimulus, an alternative explanation for the difference in decoding accuracy between the first and third stimulus is that the third stimulus was more negatively impacted by the encoding and maintenance of the target stimulus than the first.

An asymmetry in the masking effects may explain the difference in precision of observers’ reproduction responses. Without the premature decline in the neural representation introduced by the onset of the third stimulus in sequences with 50 ms ISIs, the later processing of the second stimulus remains undisturbed. This later processing may include the encoding to an observers visual working memory (Polich, 2007). If this entry to working memory is disrupted or the encoded item is degraded, the representation retrieved at reproduction will be less precise.

Our findings suggest that forward masking may be due to interference caused by reactivation of the representation of the previous stimulus, rather than recurrent processing limits (Lamme & Roelfsema, 2000; Tapia & Beck, 2014). Forward masking was thought to be due to the later processing of the masking stimulus interrupting the early processing of the masked stimulus (Keysers & Perrett, 2002).

The time course of the neural representation observed here provides an alternative explanation. When the succeeding stimulus’ neural representation peaks in decoding accuracy, the existing neural representation of the preceding stimulus decreases in accuracy. This suggests interference produced by the incoming stimulus, which is consistent with backward masking (Keysers & Perrett, 2002). When the first peak in decoding accuracy declines for the incoming stimulus, the existing neural representation seems to briefly recover before eventually tapering off. These two events appear for both ISIs and exclusively when forward masking is possible (first to second and second to third).

The brief recovery of decoding accuracy following presentation of a subsequent stimulus is reminiscent of the reactivation of activity-silent states, where weightings of synapses are primed in a configuration consistent with the activity pattern of the previous stimulus (Kamiński & Rutishauser, 2020; Oberauer & Awh, 2022; Stokes, 2015; Trübutschek et al., 2019; Wolff et al., 2017). The activity-silent state is activated by the response to the subsequent stimulus and produces a pattern that can be decoded as the previous neural representation. Unlike studies that “pinged” these activity-silent states (Wolff et al., 2017), which used tasks that demanded attention to the stimulus, the current study found similar reactivation for both task relevant and irrelevant stimuli. This may be explained by differences in temporal dynamics. That is, our stimuli could still be reliably decoded when the next stimulus was presented, whereas previous working memory experiments employed a longer delay period and pinged stimuli once they could no longer be reliably decoded (Wolff et al., 2017). Alternatively, the results may be explained by trace activity. Instead of the activity pattern of the previous stimulus becoming “silent” and only affecting the synaptic weights, it remains in a fading state that can be strengthened by any future activity (Yiling et al., 2024). The decrease in decoding accuracy that preceded the onset of subsequent stimuli could then be explained by reduced distinctiveness of the activity pattern in the presence of the new neural representation at prominence. In either case, the reactive recovery of the previous stimulus’ neural representation may interfere with the processing of the current stimulus, producing the observed forward masking.

The inconsistency in the occurrence of backward masking provides insights into its underlying neural mechanisms. The only stimuli that were susceptible to backward masking were the first and second in each sequence. Despite this, it appears to have only occurred when the ISI was 50 ms. In the time course of decoding accuracy for all three stimuli, an event can be observed exclusively when the ISI was 650 ms. Immediately preceding the onset of subsequent stimuli, the neural representation of the first and second stimuli peaked in decoding accuracy. This peak event even occurred for the third stimulus at the same relative time point (∼ -200 ms from the onset of a hypothetical fourth stimulus). The brief peak in representational fidelity occurs prior to subsequent stimulus presentation, which indicates an anticipatory response, as opposed to the conditional occurrence of the reactive activity for the first and second stimulus. Given the timing of this anticipatory boost in the 650 ms ISI condition, it is unlikely that it occurred in the 50 ms ISI sequences, as there was insufficient time between stimulus presentations. The absence of this anticipatory event in the 50 ms ISI condition may explain the why backward masking was observed more clearly in these sequences. In particular, the event may reflect a protective mechanism that reinforces the neural representation of stimuli in expectation of new incoming information. Overall, findings of decreased decoding accuracy for the first stimulus in the 50 ms, but not 650 ms, ISI sequences is consistent with previous behavioural masking studies (Bachmann & Allik, 1976; Schiller & Smith, 1965), which show backward masking is strongest when the masking stimulus is presented 30–90 ms after the target stimulus and does not produce any effect beyond 100 ms.

When implementing RSVP, the stimulus durations and ISIs have been the targets of manipulation in mitigating potential neural interference (Potter, 2012). Our findings validate the approach of increasing ISI as it does indeed sharpen neural representations but increasing stimulus durations do not. Instead, shortening stimulus durations provides greater data collection opportunity and improves analytical power. In an additional paradigm, distinct patterns in decoding accuracy were observed for stimuli subjected to forward and backward visual masking. When backward masking was weaker, an anticipatory increase in decoding accuracy was seen whilst a reactionary increase was observed for all forward masking opportunities. From these findings, some of the proposed mechanisms of the masking effects can either be eliminated or refined (Keysers & Perrett, 2002).

A limitation of the current study is the inability to determine whether the neural events found were localised to specific cortical areas or not. While whole brain decoding provides more comprehensive estimates of neural representations, future studies could employ neuroimaging techniques with greater spatial resolution to determine whether these neural mechanisms are localised and if visual masking is a low-level phenomenon (Green et al., 2005; Yiling et al., 2024). Similarly, a paradigm involving complex stimuli (e.g., faces, naturalistic scenes, etc.) or a response task involving higher-level meaning would allow comparison of visual masking beyond the low-level spatial location masking in the current study (Potter, 2012; Potter et al., 2014; Retter et al., 2018). Another limitation of the current study is the binarization of ISI in the second experiment. Using graded ISIs would facilitate observation of whether features (onset time, duration, and strength) of the events in decoding accuracy demonstrate graded differences like peak decoding accuracy did and determine the time required for the conditional anticipatory event to occur.

## SUPPLEMENTARY MATERIALS

**Figure S1.**
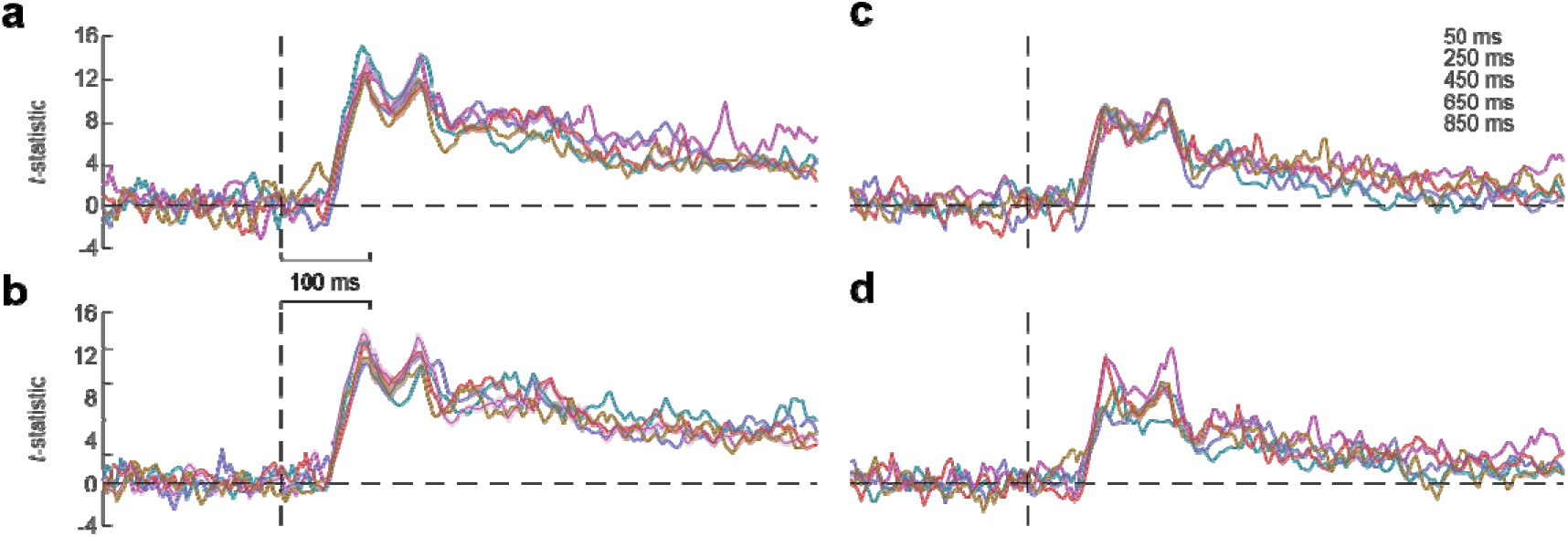
Effect size of stimulus duration and ISI on neural representations. **a**) *t*-statistics for the five stimulus durations on decoding accuracy, averaged over interstimulus interval. **b**) Same as (**a**), but for the five interstimulus intervals, averaged over stimulus duration. (**c, d**) Same as (**a, b**), but matching the number of presentations (70) across conditions. The shaded regions in (**a**) and (**b**) represent ±1 SEM, calculated from bootstrapping (*n* = 1000).

**Figure S2.**
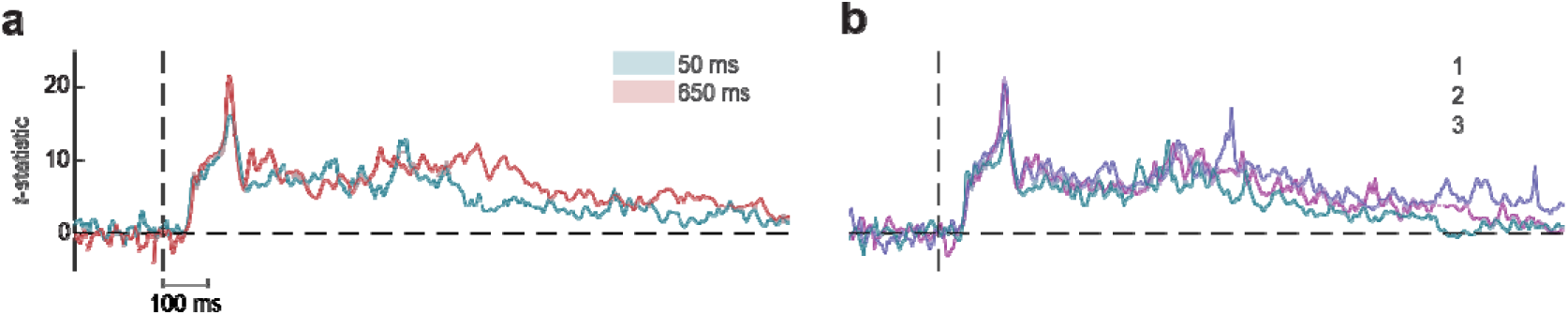
Effect size of ISI and order on neural representations. **a**) *t*-statistics for the interstimulus interval on decoding accuracy, averaged over sequence position. **b**) Same as (**a**), but for the sequence positions, averaged over ISI. The shaded regions in (**a**) and (**b**) represent ±1 SEM, calculated from bootstrapping (*n* = 1000).

